# Expansion history and environmental suitability shape effective population size in a plant invasion

**DOI:** 10.1101/416693

**Authors:** Joseph E Braasch, Brittany S Barker, Katrina M Dlugosch

## Abstract

The margins of an expanding range are predicted to be challenging environments for adaptation. Marginal populations should often experience low effective population sizes (Ne) where genetic drift is high due to demographic expansion and/or census population size is low due to unfavorable environmental conditions. Nevertheless, invasive species demonstrate increasing evidence of rapid evolution and potential adaptation to novel environments encountered during colonization, calling into question whether significant reductions in Ne are realized during range expansions in nature. Here we report one of the first empirical tests of the joint effects of expansion dynamics and environment on effective population size variation during invasive range expansion. We estimate contemporary values of Ne using rates of linkage disequilibrium among genome-wide markers within introduced populations of the highly invasive plant *Centaurea solstitialis* (yellow starthistle) in North America (California, USA), as well as in native European populations. As predicted, we find that Ne within the invasion is positively correlated with both expansion history (time since founding) and habitat quality (abiotic climate). History and environment had independent additive effects with similar effect sizes, supporting an important role of both factors in this invasion. These results support theoretical expectations for the population genetics of range expansion, though whether these processes can ultimately arrest the spread of an invasive species remains an open question.

## Introduction

Adaptation is expected to be a critical component of how species respond to novel environmental conditions, such as those encountered during colonization and range expansion (Mayr 1963; Kirkpatrick & Barton 1997; Griffith & Watson 2006; Colautti & Barrett 2013; Bock et al. 2015; Hamilton et al. 2015). At the same time, it has been suggested that colonizing species might often experience small population sizes that limit the ability of founding populations to respond to natural selection (Elam et al. 2007; Dlugosch et al. 2015b; Welles & Dlugosch 2018). Small population sizes could result from both founder events and maladaptation to novel environments. A failure to adapt under these conditions could slow or limit range expansion and contribute to the formation of range limits (Bridle & Vines 2007; Eckert et al. 2008; Sexton et al.2009). These effects are currently an active area of theoretical and experimental research (Gilbert et al. 2017; Szűcs et al. 2017a; Szűcs et al. 2017b), but there is little empirical information about the dynamics of population size and its influence on evolution during ongoing range expansions in the wild (Ramakrishnan et al. 2010; Wootton and Pfister 2015).

Population genetic models predict that adaptation can fail during range expansion due to the strong effects of genetic drift during colonization (Lehe et al. 2012; Peischl et al. 2013;Peischl et al. 2015). Range expansions are expected to involve a series of founding events,resulting in reduced effective population size (*N* _*e*_) and increased sampling effects as the range boundary advances (Le Corre & Kremer 1998; Excoffier 2004; Slatkin & Excoffier 2012). In particular, low *N*_*e*_ at the leading edge can cause random alleles, including deleterious mutations, to ‘surf’ to high frequency regardless of patterns of selection (Hallatschek et al. 2007; Excoffier and Ray 2008; Excoffier et al. 2009; Moreau et al. 2011). This can create an ‘expansion load’ of deleterious alleles at the wave front, but it is also possible that beneficial mutations can surf to high frequency and aid in local adaptation (Lehe et al. 2012; Peischl et al. 2013; Peischl et al. 2015). The effects of range expansion on adaptation have been empirically observed with greatest detail in bacterial culture, where manipulative experiments have shown that the strength of genetic drift is key to determining whether allele surfing promotes or hinders adaptation (Hallatschek & Nelson 2010; Gralka et al. 2016).

Ecological conditions should also shape *N*_*e*_ during range expansion via their impact on population (census) size and demography. If leading edge environments are novel relative to those experienced by source populations, then founding genotypes will not be pre-adapted and are likely to experience lower absolute fitness. When populations are at relatively low abundance and/or fluctuate in size due to unfavorable conditions, *N*_*e*_will be reduced relative to larger or more stable populations (Wright 1938; Crow & Morton 1955; Kimura & Crow 1963; Frankham 1996). In a rare empirical example, Micheletti & Storfer (2015) found that streamsidesalamander (*Ambystoma barbouri*) populations on the periphery of the range were also on the margins of their climatic niche and tended toward lower *N*_*e*_. Similarly, peripheral populations of North American *Arabidopsis lyrata* posess greater genetic load and appear to exist at their ecological, and perhaps evolutionary limits (Willi et al. 2018). These studies address a set of long-debated hypotheses proposing that range limits form in part because they include ecologically and/or genetically marginal populations (Kirkpatrick & Barton 1997; Phillips 2012; Chuang & Peterson 2016), which fail to support local adaptation and further expansion (i.e. the ‘central-marginal’, ‘center-periphery’ and ‘abundant center’ hypotheses: (Sagarin and Gaines 2002; Eckert et al. 2008; Pironon et al. 2015). Importantly, all the above hypotheses share the prediction that colonization will be associated with reduced response to selection for ecological reasons independent of any population genetic effects of range expansion itself. The relative importance of these two factors (ecology and expansion) for shaping *N*_*e*_ at range margins is unknown, but both have the potential to reduce opportunities for local adaptation.

Although *N*_*e*_ has long been used as a fundamental measure of the scale of genetic drift in populations (Wright 1931; Robertson 1960; Kimura and Crow 1963; Kimura 1964; Ohta 1992; Charlesworth 2009), there is surprisingly little known about how *N*_*e*_ changes during the process of range expansion. Most empirical population-level estimates come from the field of conservation genetics, where *N*_*e*_is used to infer the potential for genetic drift to exacerbate the decline of threatened populations (Lynch et al. 1995; Frankham 1996; Sung et al. 2012). These studies have demonstrated that *N*_*e*_ can be highly variable within species, sensitive to local demography and modes of reproduction, and captured only in part by measures of census size (Frankham 1995; Palstra and Ruzzante 2008). For example, in recovering Chinook salmon (*Orcorhynchus tshawytscha*) populations, Shrimpton and Heath (2003) found up to a three-fold difference in both *N*_*e*_ and its ratio with census size across spawning sites. While low *N*_*e*_is generally expected in declining populations, many of the same demographic factors are likely to affect *N*_*e*_ in founding populations (Colautti et al. 2017).

Despite the potential obstacle low *N*_*e*_ might pose to adaptation, many species -- including large numbers of invaders -- have been successful at colonization and show evidence of adaptive evolution during range expansion (Rice and Mack 1991; Dlugosch and Parker 2008; Linnen et al. 2009; Colautti and Barrett 2013; Vandepitte et al. 2014; Colautti and Lau 2015; Li et al. 2015). At the same time, detailed studies of range expansion have found some evidence of serial founding events and associated increases in genetic drift (Ramakrishnan et al. 2010; Graciá et al. 2013; White et al. 2013; Pierce et al. 2014), and it is notable that few invasions have been convincingly shown to have expanded beyond the fundamental niches of their native range (Petitpierre et al. 2012; Tingley et al. 2014). Taken together, it appears that adaptive evolution might be achievable in many invading species, but that perhaps expansion load and ecological mismatch may act, either independently or in concert, to prevent further expansion under some conditions. An understanding of how founding dynamics and marginal environments have shaped *N*_*e*_ in individual wave front populations is needed to connect theoretical expectations to these observed patterns of successful range expansion.

Here we estimate contemporary *N*_*e*_ for populations of the obligately outcrossing annual plant *Centaurea solstitialis* (yellow starthistle) across its invasion of California (USA) and its native range in Eurasia. In California, *C. solstitialis* was initially introduced in the mid 19th century into the San Francisco Bay area as a contaminant of alfalfa seed (Gerlach 1997; DiTomaso et al. 2006). By the mid 20th century, the species was rapidly expanding through California’s Central Valley and Sierra Nevada foothill grasslands, and the current leading edge of this invasion lies above 4000 m in elevation along the west side of the Sierra Nevada Mountains (Pitcairn et al. 2006). During this history of invasion, *C. solstitialis* has crossed climatic environmental gradients that are largely independent in direction from the pathway of colonization (Fig. 1), allowing us to quantify the influence of both climate and expansion history on estimates of *N*_*e*_across populations.

**Fig 1.**
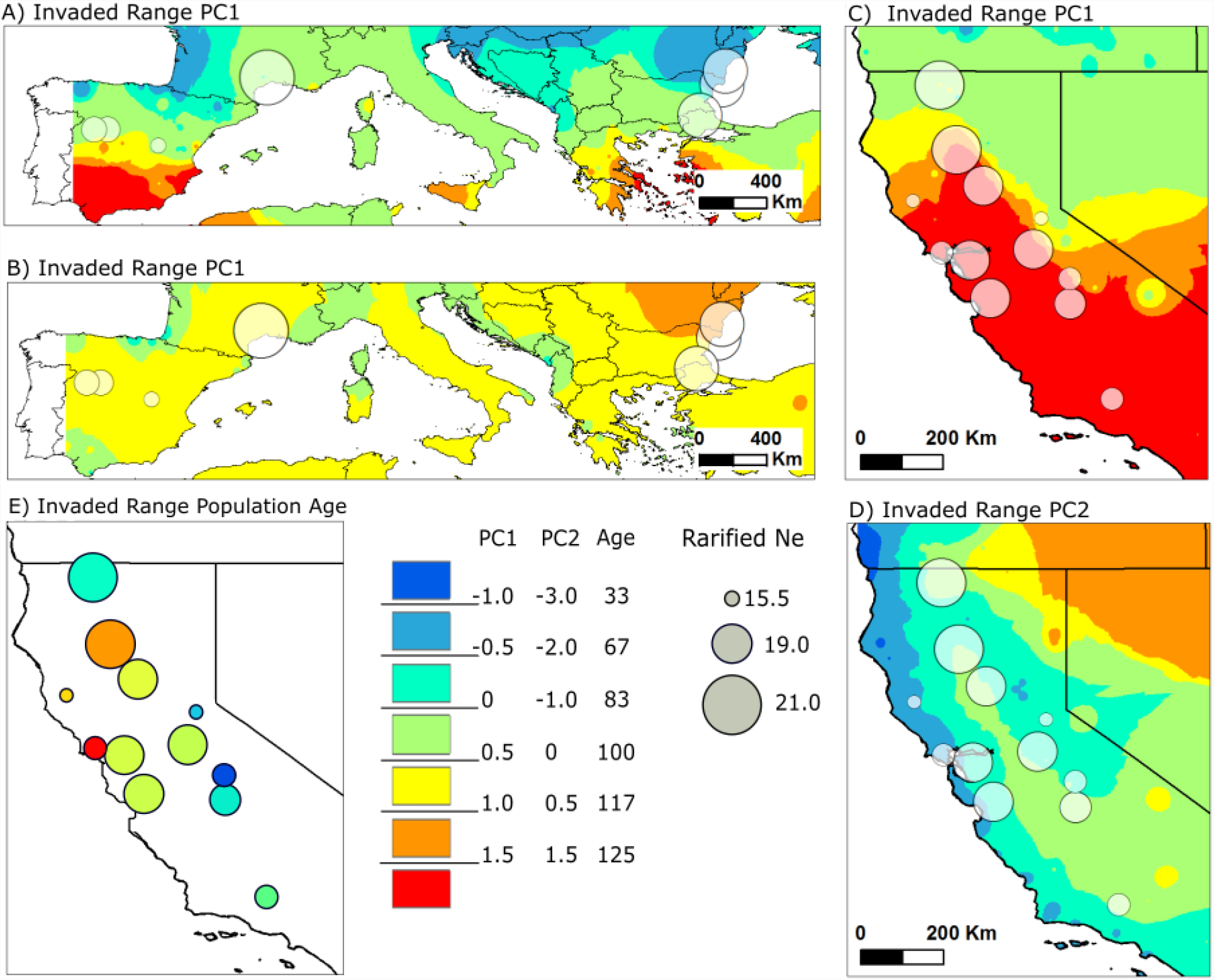
The distribution of rarefied *N*_*e*_, climatic principal component (PC) gradients, and population age across *C. solstitialis* populations in Eurasia and California. In all panels, circles indicate sampled populations with a diameter proportional to *N*_*e*_. PC1 is positively correlated with annual temperature and temperature of the driest quarter and negatively correlated with seasonal differences in total radiation in the native (a) and invaded (c) ranges. PC2 is positively correlated with seasonal differences in temperature and negatively correlated with annual precipitation and seasonal differences in precipitation in the native (b) and invaded (d) ranges. In the California invasion, population age (e) reflects a history of expansion beginning in the San Francisco Bay area and expanding first to the North and then to the South and East of the state.

We used Restriction-site Associated DNA sequencing (RADseq) to estimate contemporary *N*_*e*_in *C. solstitialis* populations sampled at a single time point. In addition to testing for the joint influence of expansion dynamics and ecological conditions on *N*_*e*_ in this system, we explored solutions for general problems associated with using large genome-wide marker data sets like this to estimate *N*_*e*_. Linkage disequilibrium *N*_*e*_ (LD-*N*_*e*_) is a powerful method for inferring contemporary *N*_*e*_ from single time sampled data, and does so by utilizing the frequency of statistical linkage across loci (Waples and Do 2008; Gilbert and Whitlock 2015). This method requires that loci segregate independently of each other, and while RADseq is widely used to produce population genetic datasets in non-model systems (Narum et al. 2013;Catchen et al. 2017), it is likely to violate this assumption of independence, resulting in biased calculations of *Ne*.

We used marker resampling and rarefaction approaches to improve inferences of variation in *N*_*e*_ across populations. We tested for effects of expansion history (time since founding) and habitat quality (climatic variables) on rarified *N*_*e*_ estimates, and compared these values to those from populations in the native range. We also explored the ability of genetic diversity measures to predict values of *N*_*e*_, given that non-equilibrium population dynamics may in the short term decouple contemporary *N*_*e*_ from its expected long term effects on genetic variation (e.g. Nei et al. 1975; Varvio et al. 1986; Alcala et al. 2013; Epps and Keyghobadi 2015). Together these analyses shed light on the factors shaping the fundamental parameters of evolution during colonization and range expansion.

## Materials and Methods

### Genomic Data

Genome-wide markers for *C. solstitialis* in this study were sampled from single nucleotide polymorphisms in double-digest RADseq (ddRADseq; (Peterson et al. 2012),previously published by Barker and colleagues (Barker et al. 2017; Dryad doi:10.5061/dryad.pf550). All sequences were obtained from *C. solstitialis* individuals germinated in the laboratory from wild collected seed. Seeds were sampled in 2008 from maternal plants along a linear transect in each population, with >1m separation between individuals. Populations included at least 13 individuals each grown from different maternal plants, from 12 invading populations in California and 7 native populations in Europe (451 individuals total; Table S1).

Briefly, sequence data published by Barker and colleagues (2017) were generated as follows. Genomic DNA was extracted with a modified CTAB protocol (Webb and Knapp 1990) and fragmented using *Pst*I and *Mse1* restriction enzymes. Samples were individually barcoded, cleaned and size selected for fragments between 350 and 650 bp. Size selected fragments were amplified through 12 PCR cycles and sequenced on an Illumina HiSeq 2000 or 2500 platform (Illumina, Inc., San Diego, CA USA) to generate 100 bp paired-end reads. Reads were de-multiplexed with custom scripts and cleaned with the package SNOWHITE 2.0.2 (Dlugosch et al. 2013) to remove primer and adapter contaminants. Barcode and enzyme recognition sequences were removed from individual reads, and bases with phred quality scores below 20 were clipped from the 3’ end. Reads were trimmed to a uniform length of 76 base pairs. The R2 (reverse) reads from the data set were removed due to variable quality, and all analyses in this study were conducted using R1 (forward) reads only.

We used the denovo_map.pl pipeline in STACKS 1.20 (Catchen et al. 2011; Hohenlohe et al. 2011) to identify putative alleles within individuals, allowing a maximum of two nucleotide polymorphisms when merging reads (-M parameter in STACKS) a maximum of two alleles per locus (-X), and a minimum coverage depth of 5 (-m). A catalog of loci and single nucleotide polymorphisms (SNPs) was generated across individuals, allowing two polymorphisms (-n) between individuals within a stack. The population.pl module in STACKS was used to calculate the population level nucleotide diversity (π) (Nei & Li 1979; Allendorf 1986). We restricted our analyses to those loci that were sequenced in 80% of individuals within a population and 90% of all populations (-r and -p parameters respectively).

### Estimates of Ne

We used SNPs identified by STACKs to calculate *N*_*e*_for each population using a method based on linkage disequilibrium among loci with a correction for missing data (Waples & Do 2008) implemented in the program NeEstimator v.2.01 (Do et al. 2014). This method derives estimates of *N*_*e*_ from the frequency of statistical linkage among loci and has been shown to be one of the best predictors of *N*_*e*_ (hereafter LD-*N*_*e*_) for markers sampled at a single time point (Gilbert & Whitlock 2015; Wang 2016; Waples 2016). The LD-*N*_*e*_ method is not strongly influenced by the total genetic diversity in the sample (Charlesworth 2009; Do et al. 2014), making it particularly well suited to analyses of invading populations where low genetic diversity might arise from founder effects unrelated to the number of individuals currently reproducing in the population. We used a minimum allele frequency threshold of 0.05 for including a locus in the analyses, which was the lowest threshold that did not result in excessive loss of loci and infinite estimates of *N*_*e*_ at some study sites.

The ddRADseq data yielded thousands of SNPs across the genome, 622 of which passed our screening requirements (see Results). Some of these loci were located in the same RAD 76bp sequence, and we expect that these and many others do not segregate independently in our data set, either due to physical proximity or the influence of selection on multi-locus allele combinations (*C. solstitialis* has a genome size of 850Mbp, distributed across 8 chromosomes (Bancheva & Greilhuber 2005; Widmer et al. 2007). To increase the likelihood of basing our estimates on independent loci, we re-sampled random sets of 20 polymorphic SNPs from unique sequences to obtain distributions of LD-*N*_*e*_ estimates for each population. Each population was resampled 30,000 times. Sampling distributions were generally lognormal (Supporting Information Fig. S1), and we used medians to identify the peak estimate.

We observed a strong, positive effect of the number of individuals sampled in each population on median LD-*N*_*e*_ (F_1,17_=9.36, P=0.007). Unequal sampling has been shown to decrease the accuracy of LD-*N*_*e*_ estimates (England et al. 2006; Authors), and NeEstimator implements a corrective algorithm to address this problem (Do et al. 2014). To account for persistent sampling effects we produced rarefaction curves of median *N*_*e*_estimated by subsampling different numbers of individuals (10 to the maximum number available per population) after marker resampling. As above, each marker resampling consisted of 30,000 N_e_ estimates with 20 loci. Median estimates did not asymptote at our maximum sampling effort and increased linearly (see Results). We fit a linear mixed model with random intercept and slopes implemented in the Lme4 package in R (Bates et al. 2014) to obtain population specific functions which describe the relationships between the number of individuals sampled in each population and median LD-*N*_*e*_ values. The estimated slope and intercepts for each population were extracted from the model and used to calculate rarefied *N*_*e*_ for each population at a standard value of 10 individuals. We explored the relationship between rarified *N*_*e*_ and measures of genome-wide marker variation using nucleotide diversity (π) at variable sites, as calculated in STACKS. We used linear regression to predict nucleotide diversity from N_e_within geographic region (invading California populations or native European populations).

### Effects of Expansion History and Environment on N_e_

We tested for an effect of population age since founding on the rarified *N*_*e*_ of invading populations. We estimated the date of colonization for each population by searching the Jepson Online Herbarium (http://ucjeps.berkeley.edu/) for records of *C. solstitialis* in California since its first record in 1869. For each sampling location, we used the earliest date on record for the county, or for an adjacent county when the sampling location was closer to older collection records there. These dates were subtracted from the year of our seed collections (2008) to produce values of population age used in subsequent analyses. Using herbaria records to assign population ages in this manner may not represent the true population age because of the time between population founding and the first records of the population. Nevertheless, *C. solstitialis* has a relatively well-documented invasion history in California (858 specimens on record, 577 records with GPS data, 61 records prior to 1930 in the Jepson Herbarium), and our population age estimates are in line with historical reconstructions of a general pattern of expansion out from initial establishment in the San Francisco Bay area first to the Central Valley and then to the North, East, and South (Gerlach 1997; DiTomaso et al. 2006; Pitcairn et al. 2006).

We also tested for the influence of the climatic environment on rarefied *N*_*e*_ in both invading and native populations. To quantify the climatic gradients that might be most relevant to *C. solstitialis* ecology, we used the first two principal component (PC) axes of climatic variation across *C. solstitialis* collection sites in North America and Europe, as previously identified by Dlugosch and colleagues (2015a; Fig. 1). Climond variables at 18.5 × 18.5 km resolution (Kriticos et al. 2012) were extracted for each site using ArcGIS. Larger values along the first PC climate axis generally indicate sites with higher temperatures and lower seasonality in total radiation. Larger PC2 values indicate lower annual precipitation and greater seasonality in temperature. Greater seasonal variation in temperature has been shown to be related to ecologically important traits (plant size and drought tolerance) in *C. solstitialis* in both the native and invaded ranges (Dlugosch et al. 2015a).

To quantify the contributions of both population age and climatic environment to variation in rarified *N*_*e*_ for the invaded range, we used a general linear model with *N*_*e*_ as the dependent variable and population age, climate PC1, climate PC2, and their interactions as explanatory variables. We constructed a separate model of rarified *N*_*e*_ in native range populations using only PC1 and PC2 as variables, since no information about population age is available for the native range. We used model decomposition and F-scores to identify the best fit model. To explore the relative effect of each variable and their interactions on *N*_*e*_, effect sizes were calculated as partial eta-squared values, which partition the total variance in a dependent variable among all independent variables (analogous to R^2^ in multiple regression), using the best fit linear model with the function ‘etasq’ in the R package ‘heplots’ version 1.3-1 (Fox et al. 2008). We also tested for an overall difference in the rarified *N*_*e*_ of invading and native populations using a Wilcoxon signed-rank test and a Monte Carlo exact test implemented with the coin package in R (Hothorn et al. 2006).

## Results

We obtained a dataset of 622 SNPs from our filtering pipeline which were used to estimate *N*_*e*_. For each population, subsampling produced a wide range of estimates of LD-*N*_*e*_, contingent upon which 20 loci were sampled (Supporting Information Fig. S1). These distributions spanned at least 4 orders of magnitude for every population. Distributions peaked strongly around median estimates (Supporting Information Fig. S1). Median estimates of LD-*N*_*e*_ prior to rarefaction varied from 19.5 to 38.5 across the California invasion (Supporting Information Table S1). In general, estimates were higher in central and northern California and decreased to the East and South (Fig. 1). In native populations, median *N*_*e*_estimates ranged from 16.2 to 42.7, with the three populations with lowest LD-*N*_*e*_ located on the western side of the range in Spain (Fig. 1).

We found a strong association between median LD-*N*_*e*_ and the number of individuals sampled from a population (r^2^_adj_ = 0.32, F_1,17_= 9.36, P=0.007). Rarefaction sampling produced positive relationships between LD-*N*_*e*_ and the number of individuals resampled within each population (Fig. 2). Slopes ranged from ~0.11 to 1.19. Importantly, rarefaction removed the significant effect of sampling effort on *N*_*e*_ values (rarefied Ne vs. total sample size; r^2^_adj_ = .12, F_1,17_=3.47, P=0.08).

**Fig 2.**
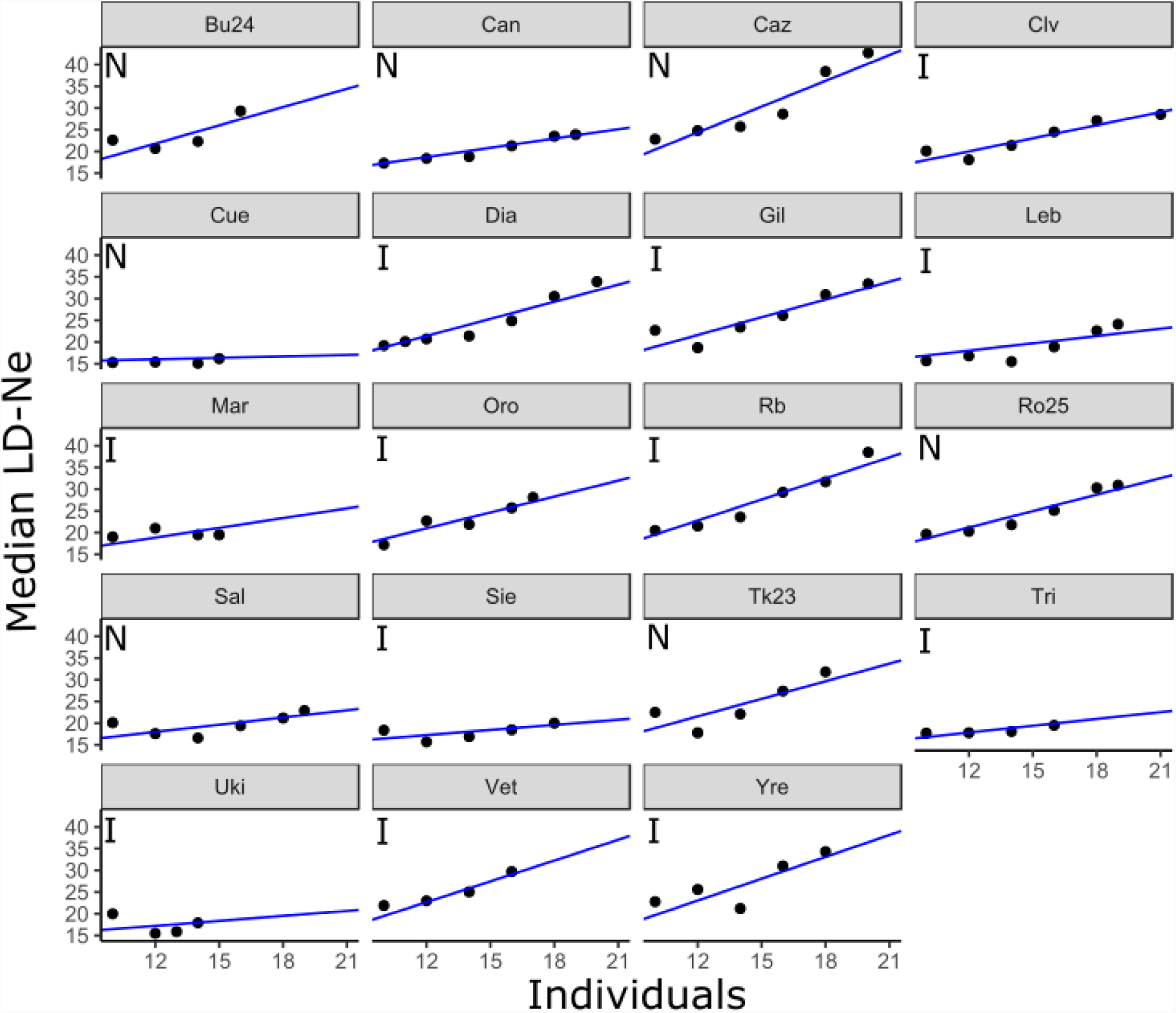
Rarified relationships for the number of subsampled individuals and median values of LD-*N*_*e*_. Rarefaction was performed by linear mixed model with random slopes and intercepts. N-Native populations; I-Invading populations.

Both climate and population age predicted rarified *N*_*e*_ in invading populations. The best fit linear model (r^2^_adj_ = 0.46, F_(4,8)_=4.09, *P*=0.0493) included significant, additive effects of population age and PC2 (Fig. 3; Table 1). Population age and PC2 were both positively correlated with rarified *N*_*e*_ values, indicating that *N*_*e*_ is largest in older populations and habitats with more temperature seasonality and lower precipitation. PC2 had a greater influence on rarified *N*_*e*_ values than age, based its larger effect size (Table 1), although this difference was small. In contrast, rarified *N*_*e*_ of native range populations was not predicted by either climatic PC variable (Full model: r^2^_adj_ = 0.2314, F_2,4_ =1.90, P=0.23) (Interactions: PC1: P=0.13, PC2: P=0.20).

**Table 1.** Individual effects for the best fit linear model explaining rarefied effective population size (*N*_*e*_) in invading populations of *C. solstitialis*, as a function of climatic principal component variables (PC1, PC2) and population age (Age).

| Effect | Coefficient | Standard Error | t-value | p-value | Effect Size |
| --- | --- | --- | --- | --- | --- |
| PC1 | 1.45 | 0.483 | -0.66 | 0.528 | 0.052 |
| PC2 | -0.39 | 0.438 | 2.50 | 0.011 | 0.579 |
| Age | 0.03 | 0.012 | 3.31 | 0.037 | 0.439 |

**Fig 3.**
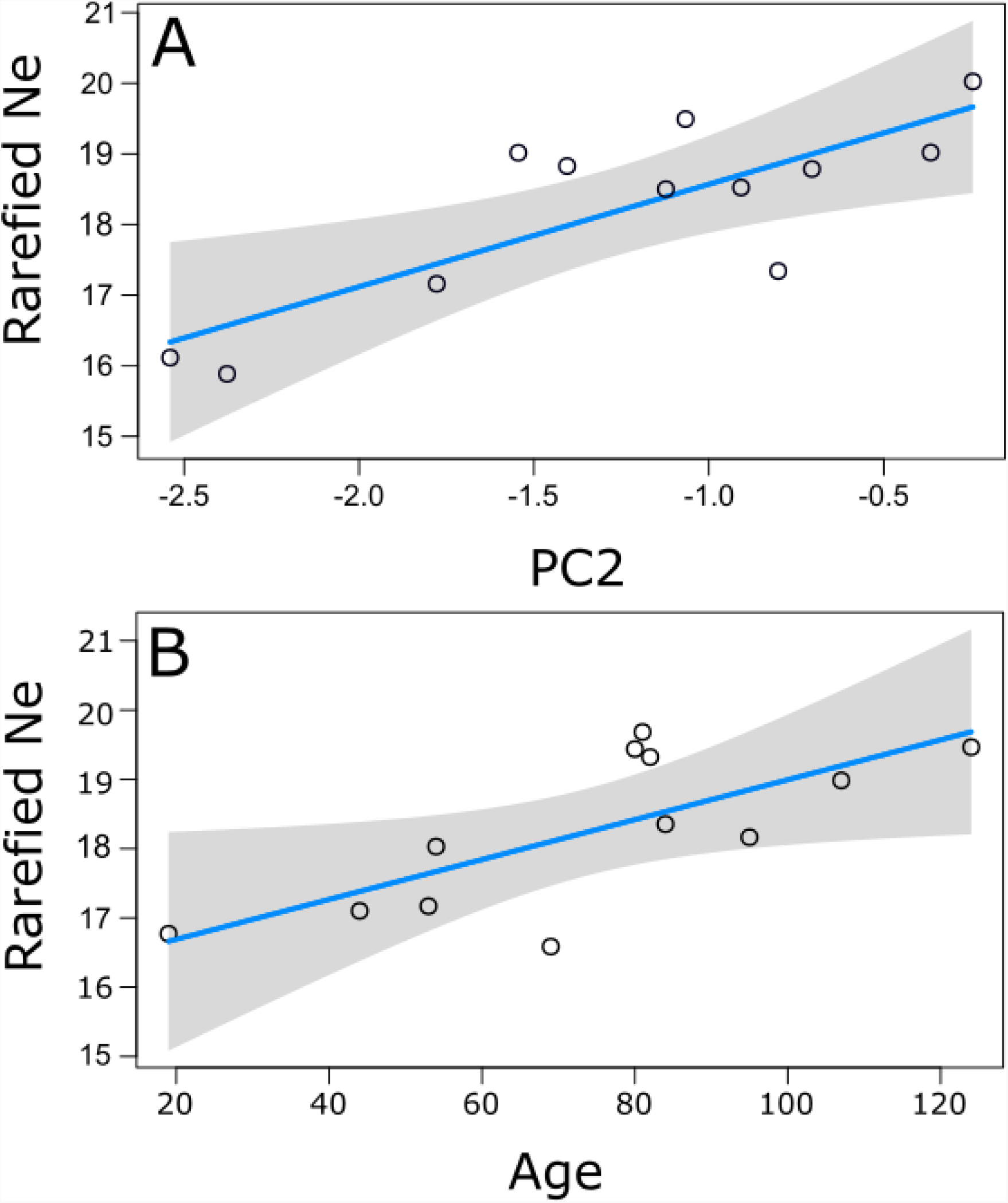
Rarefied *N*_*e*_ values are predicted by climatic variables and population age in invading C. solstitialis populations. The second principal component (PC2) of climatic variability, for which larger values represent lower annual precipitation and greater seasonality in temperature, is significantly correlated with rarefied *N*_*e*_ (A; P=0.011) as is estimated population age (B; P = 0.037). Lines show linear model fits and shading indicates the 95% confidence interval. Points represent partial residuals after taking into account other variables in the linear model.

Invading populations included a narrower range of rarified *N*_*e*_ values, nested within the distribution observed for native populations (Supporting Information Fig S2), and there was no significant difference between rarified *N*_*e*_ values in the native and invading ranges (Wilcoxon signed rank test: W = 43, *P* = 0.97; Monte-Carlo one-way exact test: *P* = 0.93). Nucleotide diversity (π) also did not differ overall between the native and invaded ranges (Model: r^2^_adj_ = 0.0.04, F_3,15_ =0.78, *P*=0.55; region term: t_2,15_ =-1.5, P=0.15). There was no significant relationship between π and *N*_*e*_ in invaded range populations (r^2^_adj_ = 0.037, F_1,10_ =0.59, *P*=0.46) and there was a positive, marginally significant relationship between nucleotide diversity and *N*_*e*_ in native range populations (r^2^_adj_ = 0.043, F_1,5_ =5.45, *P*=0.067), despite a smaller sample size in this range (Supporting Information Fig S3).

## Discussion

Here we report some of the first empirical evidence for the joint effects of both range expansion and climatic environment on contemporary *N*_*e*_ in natural populations. We produced rarified estimates of LD-*N*_*e*_ across 12 populations in the invaded range of *C. solstitialis* and found a significant positive relationship between population age and *N*_*e*_, a finding in line with theoretical expectations for the population genetics of expanding populations (Hallatschek et al. 2007; Excoffier & Ray 2008; Excoffier et al. 2009; Moreau et al. 2011; Lehe et al. 2012; Peischl et al. 2013; Peischl et al. 2015). We also find evidence that spatial variation in ecological conditions, specifically climate in our study, has a significant impact on *N*_*e*_ which is independent of population age. These effects of range expansion and climate were similar in magnitude, suggesting that both of these factors are likely to be important for shaping evolutionary outcomes in invading populations.

We emphasize that our rarified LD-*N*_*e*_ values do not reflect a ‘true’ *N*_*e*_ value for the populations in our study. Rather, rarified estimates here represent *relative* values of *N*_*e*_, and are thus useful for comparisons among populations. We expect asymptotic LD-*N*_*e*_ values for these populations to be larger, because we observed no asymptote with rarefaction for any of the populations in our study. Despite this, our estimates are similar to values reported in other plant and animal species using the same approach (e.g. Shrimpton & Heath 2003; Coyer et al. 2008; Wang et al. 2013; álvarez et al. 2015). The LD-*N*_*e*_ estimation method itself also has a tendency to underestimate known values of *N*_*e*_ in simulations (Gilbert & Whitlock 2015)

We found that subsampling of loci was important for using a genome-wide marker dataset to estimate LD-*N*_*e*_. Simulation studies of LD-*N*_*e*_ typically use on the order of 20 loci (England et al. 2006; Waples 2006; Waples & Do 2008; Gilbert & Whitlock 2015). Our resampling of 20 loci revealed that *N*_*e*_ estimates in *C. solstitialis* vary by at least 4 orders of magnitude when different sets of loci are used. This variation is expected given that particular sets of loci will capture different effects of physical linkage, history of selection, and chance sampling effects (Daly et al. 2001; Remington et al. 2001; Flint-Garcia et al. 2003). Resampling allowed us to leverage many combinations of loci across the genome to identify a well defined peak in the distribution of estimates. A resampling approach is likely to be generally useful for RAD-seq and other popular genome wide marker methods, particularly where a complete reference genome is not available to control for the physical arrangement of loci.

After accounting for these methodological issues, we found a significant effect of population age on differences in *N*_*e*_ across populations. Rarefied *N*_*e*_ estimates were lower in younger populations, which fits with expectations that a subset of individuals will contribute to range expansion (Excoffier & Ray 2008) and that genetic drift will be larger at the leading edge (Lehe et al. 2012; Peischl et al. 2013; Peischl et al. 2015). Estimates of contemporary *N*_*e*_ from our invading populations are within the distribution that we observe from native populations, suggesting that this species did not experience a large initial genetic bottleneck during introduction to the New World, nor exceptionally low *N*_*e*_ during range expansion (relative to values observed in native populations). This lack of evidence for a strong genetic bottleneck is inline with models of historical demography by Barker and colleagues (2017), who inferred little reduction in population size and maintenance of genetic diversity during the colonization of the Americas by *C. solstitialis*. In general, introduced species often lack strong genetic bottlenecks (Dlugosch & Parker 2008; Uller & Leimu 2011; Dlugosch et al. 2015b), and our results here demonstrate that species which avoid genetic bottlenecks during introduction may still experience declines in *N*_*e*_ at the leading edge of range expansion. Importantly, invading populations of *C. solstitialis* are an order of magnitude higher in density than native populations (Uygur et al. 2004; Andonian et al. 2011), indicating that the fraction of the census population that is contributing to evolutionary dynamics in the invasion is much lower than in the native range.

We also observed an independent positive relationship between climatic PC2 and *N*_*e*_ of invading *C. solstitialis* populations, consistent with an impact of habitat suitability on *N*_*e*_. High PC2 values reflect greater variation in seasonal temperatures and lower total annual precipitation, which typify areas of especially high *C. solstitialis* density in California (Dlugosch et al. 2015a). Previous studies in this system have proposed that *C. solstitialis* success stems from a lack effective competitors in more drought prone habitats (Dlugosch et al. 2015a), due in part to the extensive conversion of these habitats to rangeland (Menke 1989, (Stromberg & Griffin 1996)). These patterns of *C. solstitialis* density can be counterintuitive, as studies within the California invasion have found that water availability (both naturally occurring and experimentally manipulated) is strongly and positively correlated with *C. solstitialis* density and fecundity (Enloe et al. 2004; Morghan & Rice 2006; Hulvey & Zavaleta 2012; Eskelinen & Harrison 2014), suggesting that fitness should be highest in wetter areas. Our results are most consistent with the landscape pattern of abundant *C. solstitialis* in drier areas, and might therefore reflect differences in human land use and the availability of native competitors across the invaded range. An underlying relationship between *N*_*e*_ and land use in the invasion could also explain why we did not find a relationship with climate in the native range, where other features might shape habitat suitability for this species.

These differences in rarified *N*_*e*_ among invading populations were not predicted by sequence diversity (π). Nonequilibrium populations such as the invasions here are unlikely to have had sufficient time to reach equilibrium diversity at a given *N*_*e*_, and will also have been changing in size over time (Nei et al. 1975; Alcala et al. 2013; Epps & Keyghobadi 2015). Notably, we did find a marginally significant positive relationship between π and *N*_*e*_ in native range populations, which will have had more time to stabilize in population size and reach mutation-drift equilibrium. Moreover, rare alleles contribute important equilibrium genetic variation (Luikart et al. 1998) and native *C. solstitialis* populations have been previously shown to harbor more rare alleles than invading populations in North America (Barker et al. 2017). There is also a tendency for RAD-seq to underestimate π in more diverse genomes (Arnold et al.2013; Cariou et al. 2016), although given the loss of rare alleles from invading populations, we might expect this to affect native populations more strongly than invading populations.

Our results support the prediction that both range expansion and habitat quality can increase the genetic drift experienced by leading edge populations. There is particular interest in whether these effects can hinder adaptation, slow further colonization, and establish static range boundaries (Bosshard et al. 2017; Lehe et al. 2012; Peischl et al. 2013; Peischl et al. 2015; Marculis et al. 2017; Birzu et al. 2018). Recent studies have demonstrated a link between differences in historical values of *N*_*e*_ and differences in efficacy of selection across species (e.g. (Slotte et al. 2010; Jensen & Bachtrog 2011; Strasburg et al. 2011), and both theoretical and experimental studies of bacteria have shown that the process of range expansion can reduce contemporary *N*_*e*_ and impose limits to adaptation and further colonization at the expansion front (Hallatschek & Nelson 2010; Lehe et al. 2012; Gralka et al. 2016; Peischl et al. 2013; Peischl et al. 2015). Natural populations of *Arabidopsis lyrata* have been shown to demonstrate some of these effects, with greater genetic load in range edge populations associated with a lack of adaptation along an environmental cline (Willi et al. 2018). Limits to range expansion are expected to be sensitive to the specifics of evolutionary parameters in natural populations, including the magnitudes of *N*_*e*_ and selection, the amount and scale of gene flow across the expansion, and the genetic architecture of adaptive variation (Hallatschek & Nelson 2010; Lehe et al. 2012; Peischl et al. 2013; Peischl et al. 2015; Gralka et al. 2016).

The expansion ecology of *C. solstitialis* does not support the existence of maladapted edge populations. Populations of *C. solstitialis* closer to the range edge can achieve greater density than more interior populations in some cases (Swope et al. 2017), which runs counter to any expectations of high genetic load. Additionally, evolution of increased growth and earlier flowering appears to be enhancing the invasiveness of *C. solstitialis* (Dlugosch et al. 2015a), *suggesting that reduced Ne* at the range edge has not created a barrier to adaptation and further expansion. On the other hand, in areas where we would expect adaptation to be particularly critical for colonization, such as stressful serpentine soil habitats in California, *C. solstitialis* is notably rare or absent (Dukes 2002; Gelbard & Harrison 2003), though this might simply reflect the absence of relevant adaptive genetic variation. The availability of adaptive variation and the degree to which this is a limiting factor in species invasions is an active area of debate (Ellstrand & Schierenbeck 2000; Rius & Darling 2014; Bock et al. 2015), and should be particularly relevant to the colonization of habitats requiring significant niche evolution. The results reported here emphasize that an understanding of the evolutionary mechanisms that generate boundaries to range expansion in natural populations will require evaluating evidence not only for availability of adaptive variation (Dlugosch et al. 2015a), but also for an effective response to selection.

## Acknowledgements

We thank # reviewers for helpful comments on previous versions of this manuscript, and MS Barker for computational assistance. This study was supported by a USDA ELI Fellowship #2017-67011-26034 to JEB, the National Institute of General Medical Sciences of the NIH under Award #K12GM000708 through the University of Arizona Center for Insect Science to BSB, and USDA grant #2015-67013-23000 to KMD. The content is solely the responsibility of the authors and does not necessarily represent the official views of the National Institutes of Health. All authors declare no conflict of interests.

## Data Accessibility

All data files to be made accessible online (e.g. via Dryad)

## Author Contributions

JB and KMD conceived the study; JB and BSB performed the analyses; JB, BSB, and KMD wrote the paper.

## SUPPORTING INFORMATION

Supporting information is included as a single PDF document

